# Differential Network Analysis of Longitudinal Gene Expression in Response to Perturbations

**DOI:** 10.1101/2021.09.21.461277

**Authors:** Shuyue Xue, Lavida R.K. Rogers, Minzhang Zheng, Jin He, Carlo Piermarocchi, George I. Mias

## Abstract

Understanding changes in gene expression under the effects of a perturbation is a key goal of systems biology. A powerful approach to address this goal uses gene networks and describes the perturbation’s effects as a rewiring of each gene’s connections. This approach is known as differential network (DN) analysis. Here, we used DNs to analyze RNA-sequencing time series datasets, focusing on expression changes: (i) In the saliva of a human subject after vaccination with a pneumococcal vaccine (PPSV23), and (ii) in B cells treated *ex vivo* with a monoclonal antibody drug (Rituximab). Using network community detection, we revealed the collective behavior of clusters of genes, and detected communities of genes based on their longitudinal behavior, and corresponding pathway activations. We identified biological pathways consistent with the mechanism of action of the vaccine and with Rituximab’s targets. The approach may be useful in drug development by providing an effective analysis of expressing changes in response to a drug.

## 1. Introduction

Recent advances in RNA-sequencing (RNA-seq) technologies have enabled researchers to incorporate time in the analysis of large-scale gene expression data from biological systems. These time course (TC) experiments can generate longitudinal RNA-seq data as time series, with time-steps ranging from hours to weeks. This not only permits capturing the transcriptome’s dynamic regulation over time, but also generates large amounts of expression data that need to be analyzed and interpreted. While TC experiments can measure time-dependent gene expression changes, a good data analysis scheme is key for translating the experimental observations into meaningful information about the underlying biological dynamics [1, 2, 3, 4, 5, 6].

There are currently a wide variety of well-established data analysis methods for RNA-seq data. However, this is not the case for TC RNA-seq, as in this case another dimension, time, is present. Some methods originally developed for microarray sequencing have been adapted to RNA-seq time series data [3, 6, 7, 8]. Network-based analysis and, in particular, Differential Network (DN) analysis methods, have been shown to be very useful in the analysis of the dynamics of gene expression under the effect of an external perturbation [9] and could provide insightful interpretations of the RNA-seq data. DN analysis is a method based on the subtraction of one network from another, and has been adopted in many genomics studies in the past decade [10, 11, 12]. In a gene-gene correlation network (co-expression network), vertices represent genes while edges represent the correlation coefficient of the expression of two genes. Typically, these co-expression networks are weighted by the strength and the sign of the correlation between two genes. The DN analysis method uses a pairwise cancellation of nodes and edges common to two networks that describe the expression data before and after a given perturbation. In doing so, the process leaves behind interaction variations that describe the network rewiring induced by the perturbation. For instance, in gene expression studies, the DN analysis method successfully separated gene expressions under specific drug responses from generic stress responses [13]. It also aided researchers in investigating dysfunctional regulatory networks in unhealthy states, providing insights into the genetic basis of diseases [14]. By focusing on the structural difference between two networks, the DN analysis method has demonstrated its effectiveness in identifying biological activities in different states. In addition, this graph-based model offers an advantage in representing the architecture of a gene network’s overall changes where the emphasis is on the nature of interactions rather than the quantitative predictions in time.

In the present study, we applied a DN approach to RNA-seq time series datasets retrieved from two longitudinal TC RNA-seq experiments: (i) The first dataset (GSE108664) was generated from saliva samples from a healthy individual before and after the administration of the Pneumococcal Polysaccharide Vaccine (PPSV23) [15]. The primary goal of this study was to gain insights on the adaptive immune responses to PPSV23 through saliva profiling. Due to its convenience in processing relative to blood samples, saliva draws much interest for diagnostics as well as health monitoring applications. Saliva analysis can produce results in a timely manner, its collection is minimally invasive, and little training is required for saliva sampling, even for non-medically trained professionals. (ii) The second dataset (GSE100441) was generated from a time course experiment on primary B cells, where one set was treated with Rituximab and another used as an untreated control. Rituximab is known for its therapeutic use in targeting B cells [16] to treat cancers such as lymphomas and leukemias. This drug has a history of safe and effective usage since 1997 [17], and the World Health Organization (WHO) place Rituximab on their list of essential medicines [18]. Rituximab binds with CD20, expressed on pre-B and mature B cells, but not on stem cells [19]. The binding causes perturbations to intracellular signaling and membrane structure [20], mediating the cell depletion. It is worthwhile to mention that the B cell pathways of Rituximab activation have been experimentally validated [21, 22, 23, 24], which facilitates the evaluation of the effectiveness of the DN method we utilize in this work. Both the saliva and primary B cell experiments involve drug-treated samples (treatment sets) and untreated samples (control sets) monitored over time.

For both datasets, we started with building gene networks, one for each of the control and the treatment sets. We used gene-gene correlations between time series signals, over 24 hours in saliva and 15 hours in B cells, to evaluate pairwise gene connections. Graphically, the time series correlation networks built from the treatment sets summarized a system-wide pathway activation due to the perturbation, whereas the networks from the controls sets acted as the baseline. Within the DN analysis framework, we subtracted the baseline network from the one obtained using the treatment data, arriving at the final differential network.

The presence of modules, also known as communities, describes a topological property of networks [25, 26, 27, 28]. One community is a group of densely connected nodes. In the context of a biological system, nodes in the same community are assumed to be close in biological functions [29, 30, 31, 32, 33]. We exploited this property of the differential network to observe fine details of gene groups affected by the perturbation. Specifically, we employed one of the most established module detection algorithms, the Louvain method [34], to identify communities in our final differential network. We then performed Reactome [35] pathway enrichment analysis on individual communities, and finally examined the corresponding heatmaps for each community.

## 2. Results

Our RNA-seq time series raw data were retrieved from the Gene Expression Omnibus database under accessions GSE108664 and GSE100441 for the saliva and B cell experiments, respectively. The study of the immune response to the PPSV23 vaccine in saliva probed the expression of a potential 84647 gene identifiers (GENCODE annotation[36]) at 24 time points [15]. The other study of drug activation by Rituximab in B cells provided a dataset for 6 time points. Since gene co-expression networks rely on correlations, our network analysis could be prone to spurious correlations, which we removed as described in the STAR Methods.

We constructed our saliva DN by subtracting the saliva network without vaccine from the network obtained using post-vaccine data. The B cell DN in response to Rituximab was generated in a similar manner. Next, we clustered the DNs into communities using the Louvain community detection method [34]. We then conducted a Reactome Enrichment Analysis [35] using PyIOmica [37], on each community to identify significant pathways and associated genes. We also visualized the heatmaps of relative gene expression as a function of time for each community. Finally, we plotted the DNs and their major individual communities. The workflow is summarized in Figure 1, and additional details are provided in the STAR Methods.

**Figure 1:**
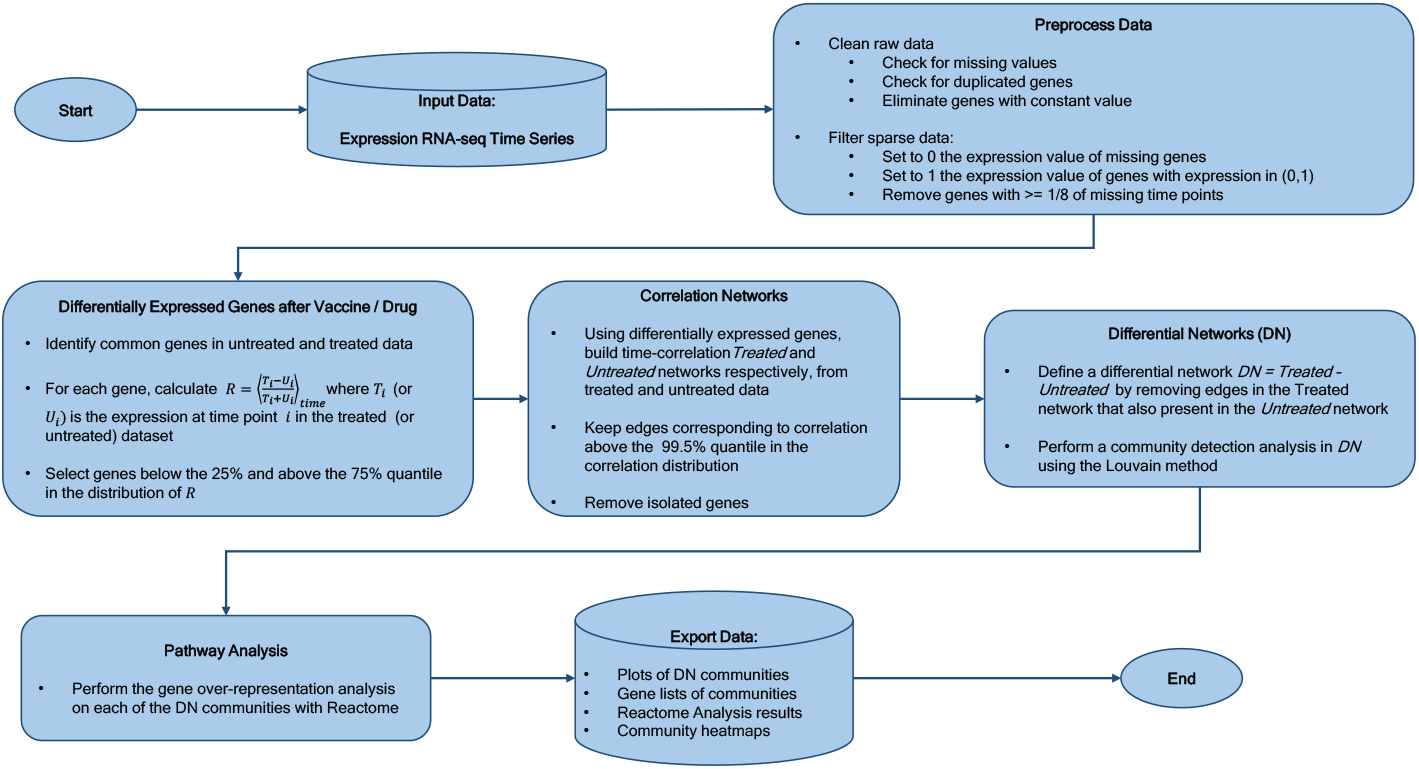
Workflow Overview. Our methodology, starts with time course experimental data, followed by network construction, differential network determination, community detection, pathway analyses of individual communities, and final results including analyses and temporal trend visualizations.

### 2.1. Saliva DN

Our saliva DN contains 1144 nodes (i.e., genes) and 13,775 edges. The Louvain algorithm identified 48 communities (modules) in total. 15 of the communities have a size of at least 4 nodes, while the remaining 33 are pairs or triplets. In the global saliva DN visualization, we excluded the communities with pairs or triplets, as none of them belonged to the three major connected components of the DN network. We also filtered the network to remove connected components with less than 4 genes. The global saliva DN is presented in Fig. 2a, where communities are visualized using different colors and encircled in loops. Furthermore, community labels are based on their size (largest to smallest, with C0 being the largest community, and C14 the smallest).

**Figure 2:**
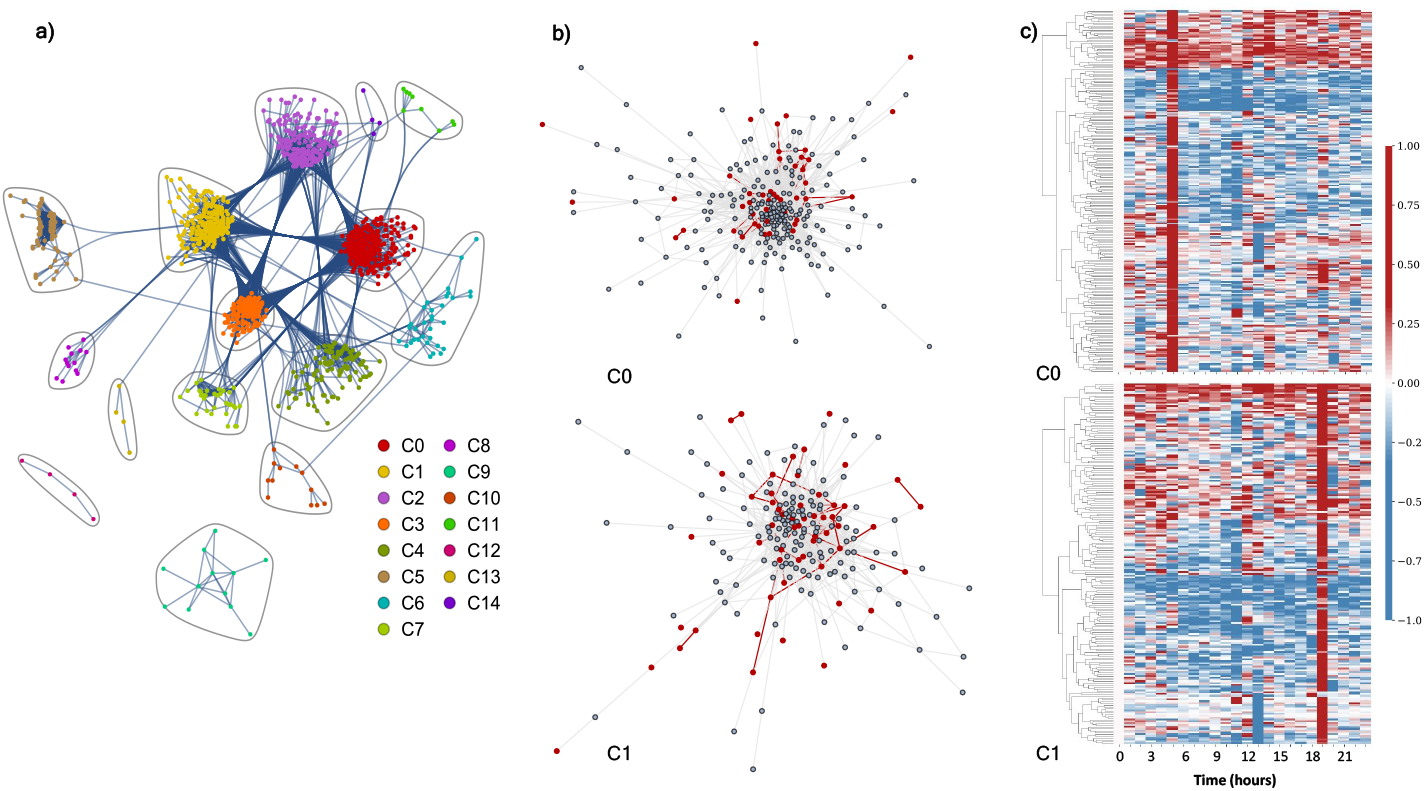
Differential network analysis for the saliva experiment. a) Differential network with community structure found by the Louvain community detection method. b) Isolated visualizations of C0 (top) and C1 (bottom) communities with red highlights corresponding to genes found in significant Reactome pathways. c) Heatmaps of C0 (top) and C1 (bottom) over 24 hours. Columns represent time points while rows denote gene identifiers. These row data demonstrate the difference in each entry’s expression relative to time 0. The relative values were determined by subtracting the individual time points from time point 0 and then normalized using a Euclidean norm, so that each row ranges from -1 (down-regulation) to 1 (up-regulation). For the dendrogram clustering we used the complete-linkage method (also known as the Farthest Point Algorithm) [55, 56].

### 2.2. Pathway Enrichment of Saliva Communities

In our pathway analysis, we queried individual communities to investigate how their highly co-expressed genes are functionally related. Our analysis is based on the Reactome pathway database [35, 38, 39].

Statistically significant enrichment of pathways (with False Discovery Rate (FDR) < 0.05) was identified in 6 communities, C0, C1, C2, C4, C8 and C9. The majority of statistically significant Reactome pathways were related to response to stimulus, immune response, and inflammatory response. Among the six communities, C0 and C1 are the two largest communities. C0 comprises of 248 genes, colored in red in the global DN shown in Fig. 2a, whereas C1 contains 198 genes, colored in yellow in the same panel. We display the C0 and C1 in Fig. 2b as representative communities. Genes that belong to the statistically significant biological pathways are highlighted in red in Fig. 2b.

In the C0 community, the Reactome enrichment analysis identified 15 statistically significant pathways (FDR < 0.05): (i) three pathways for interferon signaling, (ii) three related to the immune system, (iii) four related to antigen presentation, (iv) one associated with ER-Phagosomes, (v) one lymphoid-related, and (vi) three pertaining to interleukin-12 signaling. In particular, the alpha, beta, and gamma signaling pathways all appear in the interferon signaling pathways. The immune system pathways include one cytokine signaling and one related to the adaptive immune system. Among the four antigen-related pathways, two are explicitly associated to the dependence of Class I MHC. The Endosomal / Vacuolar pathway implies the involvement of the Class I MHC and of the Antigen processing-Cross presentation. Lastly, interleukin-12 plays a crucial role in the coordination of innate and adaptive immunity [40].

In the C1 community, the Reactome analysis identified 9 statistically significant pathways (FDR < 0.05). Two of these pathways are broadly related to the immune system and cytokine signaling. Another two pathways, the NGF-stimulated transcription and the FOXO-mediated transcription pathways, modulate cell survival, growth, and differentiation. In Table 1 we have listed all the results of the Reactome pathway enrichment analysis for C0 and C1 with FDR < 0.05.

**Table 1:**
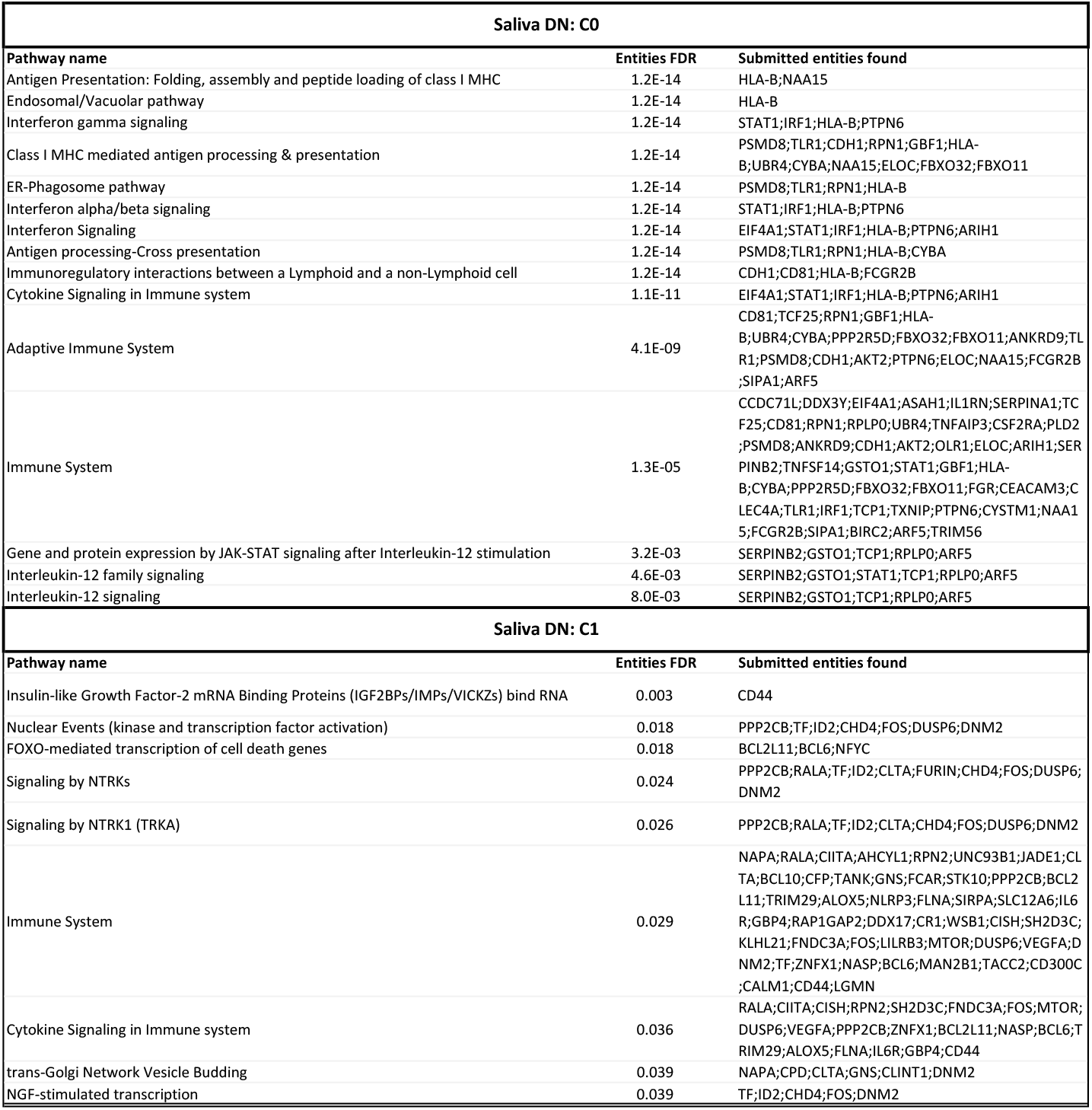
Reactome pathway enrichment analysis. Statistically significant pathways are summarized for saliva DN community C0 and C1. In the full analysis, we omitted small communities with fewer than 8 genes [57], and 12 communities (C0 to C11) qualified for the pathway analysis.

Of the communities we observed, the C0 community exhibits the strongest response to the stimulus and immune system, which is evidenced by the very low FDRs ∼ 𝒪(10^−14^). The complete pathway enrichment analysis for all communities in saliva is provided in the online data files (ODFs), available on Zenodo, in the “Results/SLV_results/reactome_analysis” folder.

### 2.3. Saliva Communities Temporal Visualization

We further visualized each community’s change over time with heatmaps within the DN network. This is shown for C0 and C1 in Fig.2c. Here, each row denotes a gene, while each column corresponds to a time point of post-treatment. The values plotted in the heatmaps are rescaled gene expression differences between the treated data and the control, and indicate the expression at the particular time point relative to the first time point of the experiment, with rows normalized using Euclidean norm. Red indicates up-regulated genes, blue down-regulated genes, and white indicates unchanged expression. The hierarchical clustering dendrograms revealed relationships among genes at each time point based on the similarity of the gene expressions. The prominent red columns show that genes are upregulated together at these time points. Note that the C0 has a pronounced peak at time point 6, making it an early responding module, while C1 is a late responding cluster, with a pronounced peak at time point 19, as illustrated in Fig. 2c.

Here we only show heatmaps for C0 and C1 as representative communities. However, we provide the other communities’ heatmaps and with their corresponding Reactome pathway analysis in the ODFs in the folder named “Results/SLV_results/network_plots”. Our saliva DN has a clear pattern of mostly discrete punctuated gene expression response times for each community. As these punctuated response times, save for one exception (both C0 and C11 show maximized response at t5), are specific to each community, they reflect the biological signatures for individual groups. Most of our saliva DN communities have only one punctuated activation time, although C5 in the saliva DN has 3 up-regulation events at time points 15, 20, and 22 that do not overlap with those of other communities. Between the communities, we observed strong temporally-specific relationships. Our heatmaps are suggestive of the presence of directional signaling between early-activation communities and subsequent groups, with a potential sequential activation pattern as follows: C6, C9, C8, C2, C0 and C1, C3, C4, C10, C5, C1, and finally C5. At time points from t6 to t10, t14, and from t16 to t18, no communities activated.

### 2.4. B Cell DN

Our B cell DN consists of 1,759 nodes (genes) and 10,421 edges that we classified into 145 communities using the Louvain algorithm. Similar to the saliva DN, most of these communities are small clusters on small components. Due to its larger size relative to the saliva DN and larger number of communities, our cutoff for plotting was increased to 8 nodes both for community and component size. The global B cell DN is presented in Fig. 3a, with 5 components and 14 communities. Here, we omitted the remaining 130 communities since they neither belong to any of the 5 main components, nor are they large enough for Reactome enrichment analysis. Like in the saliva DN, communities were ordered in descending size (largest to smallest, from C0 to C13 respectively), designated with different colors, and encircled by loops. Fig. 3 has the same format of Fig. 2. In this case, C2 and C4 are displayed in panel b, as magnified representations of the purple cluster and the green cluster, respectively, in panel a. Panel b’s magnified perspective provides details about the communities’ internal structures. In Fig 3b, for example, we observe that some of the genes highlighted in red form a clique.

**Figure 3:**
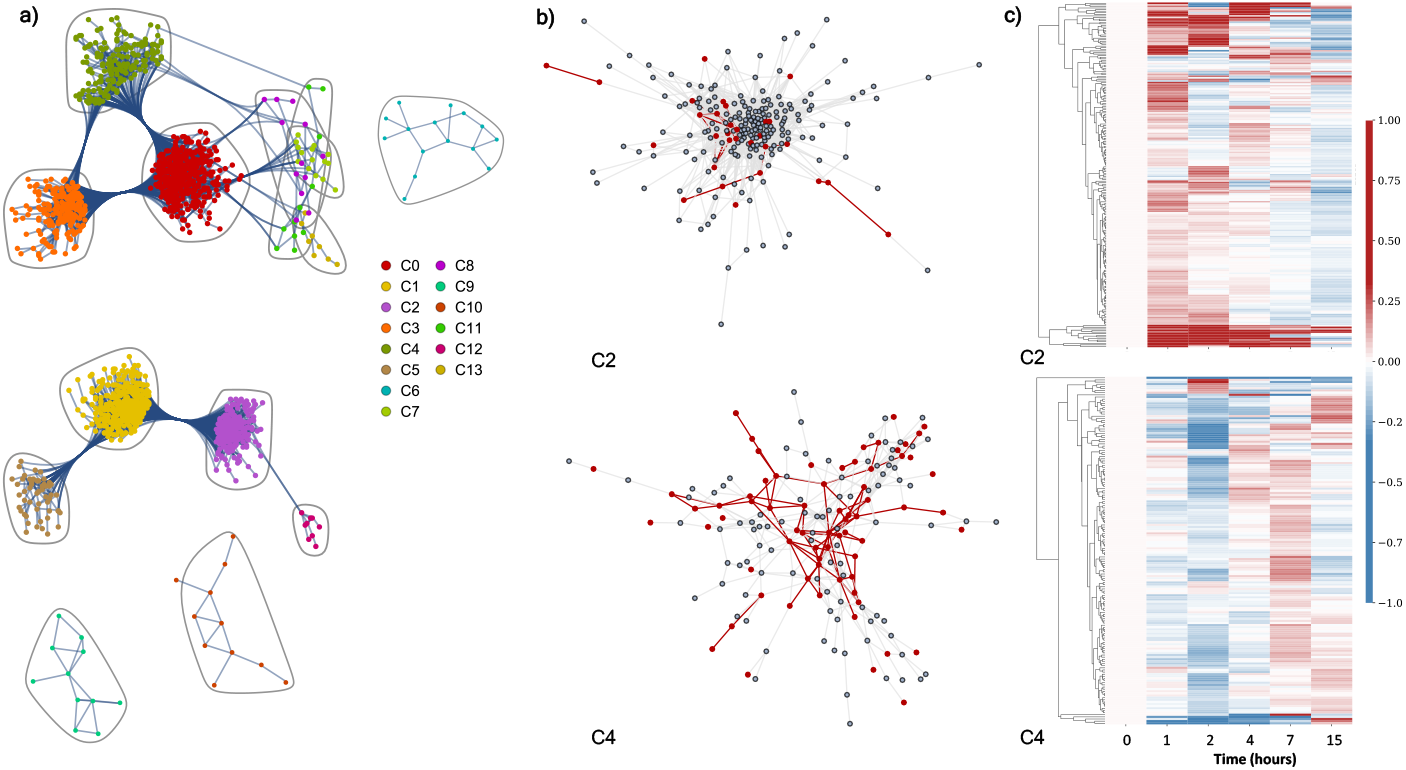
Differential network analysis for the B cell experiment. a) Differential network with community structure found by the Louvain community detection method. b) Isolated visualizations of C2 (top) and C4 (bottom) communities with red highlights corresponding to significant nodes (genes) and their edges (correlations). c) Heatmaps of C2 (top) and C4 (bottom) over 15 hours (6 time points). Columns represent time points while rows denote genes. These row data demonstrate the difference in each entry’s expression relative to time 0. The relative values were determined by subtracting the individual time points from time point 0 and then normalized using a Euclidean norm, so that each row ranges from -1 (down-regulation) to 1 (up-regulation). For the dendrogram clustering we used the complete-linkage method (Farthest Point Algorithm) [55, 56].

### 2.5. Pathway Enrichment of B Cell Communities

As for the saliva DN, we conducted a community-wise Reactome enrichment analysis for communities with at least 8 genes. 14 communities in the B cell DN were analyzed. This analysis found 9 communities with statistically significant pathway enrichment (FDR < 0.05.): C2, C4, C5, C6, C7, C9, C10, C12, and C13. Most of the pathways associated with genes in these communities centered around transcriptional regulation, protein metabolism, DNA binding ability, and signaling. Among its 111 statistically significant pathways, C4 was found to be strongly enriched with genes in the FCERI-mediated NF-*κ*B activation pathway, the B cell receptor (BCR) signaling pathway, and the Fc epsilon receptor (FCERI) signaling pathway. These pathways and others relevant to Rituximab mechanism of action, are listed in Table 2. The NF-*κ*B pathway activation by FCERI leads to the production of cytokines during mast cell activation, making it important in allergic inflammatory diseases [41]. C4 also contained a significant number of genes in the B cell receptor pathway, an important pathway related to B cells. The Fc epsilon gene is expressed on antigen-presenting cells, and its signaling occurs on the plasma membrane. A comprehensive list of statistically significant pathways can be found in the ODFs in the “Results/Bcell_results/reactome_analysis” folder.

**Table 2:**
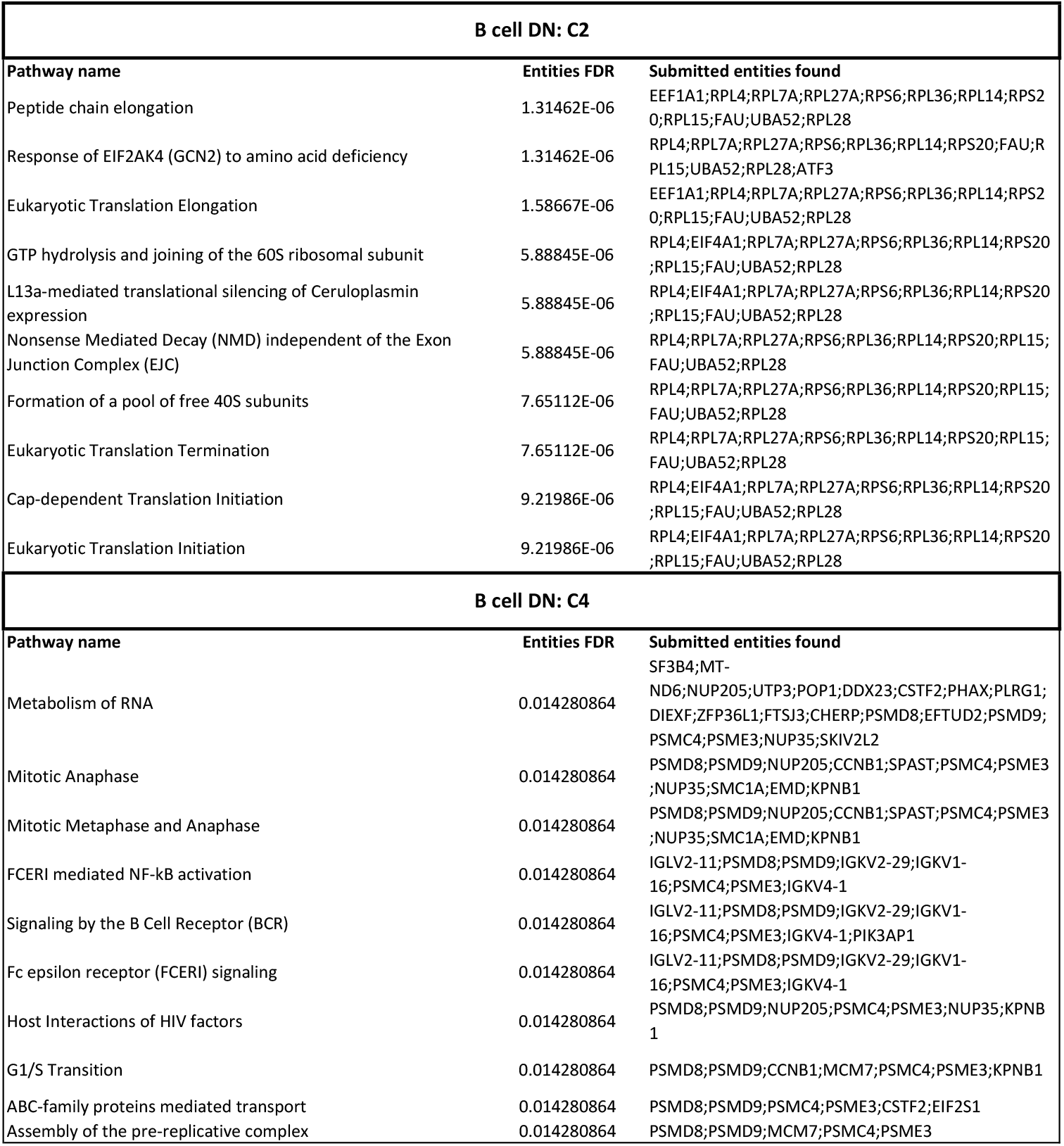
Reactome pathway enrichment analysis. Statistically significant pathways are summarized for primary B cell DN community C2 and C4. In the full analysis, we omitted small communities with fewer than 8 genes [57], and 12 communities (C0 to C11) qualified for the pathway analysis.

In summary, C4 contains the highest number of responsive pathways which are relevant to the B cell response to Rituximab. As our representative communities, we display the C2 and C4 in Fig. 3 b, our two largest among the 9 communities with significant pathways. Our top 10 pathways based on p-values from the Reactome enrichment analysis for C2 and C4 are listed in Table 2.

### 2.6. B Cell Communities Temporal Visualization

The heatmaps for the temporal behavior for the C2 and C4 communities of the B cell data are shown in Fig. 3c. The formatting of the heamaps is the same as that of the saliva heatmaps; all values in the heatmaps refer to gene expression relative to time 0 in the treated dataset. The C4’s blue column at time point 2 and the less prominent blue column for C2 at time point 15 identify patterns of down-regulation in the two community. While C2 shows a trend of initial up-regulation followed by a gradual diffusion, C4 exhibits an initial down-regulation, followed by later up-regulation.

Though C2 and C4 are our representative communities, we carried out heatmap visualization for all our 9 communities that demonstrated significant pathway enrichment. These heatmaps are available to view in the ODFs in the “Resuls/Bcell_results/network_plots/heatmaps” folder. Overall, in the B cell community heatmaps, we recognized three types of time patterns in terms of collective behavior within an individual community. In the first pattern group, the majority of genes started with a moderate degree of down-regulation. By 7 hours, most instead displayed slight or moderate up-regulation. However, each of these timepoints contained a significant minority of genes with a small level of fluctuation, with the size of the deviating group differing in each heatmap. The second observed time pattern operated in reverse, with most genes beginning upregulated and shifting towards downregulation by the 15-hour mark. Finally, a third group remained consistent in its behavior, with genes trending one way or remaining unchanged across the entire time period.

## 3. Discussion

Our goal was to determine whether the DN method can identify the activation of biological processes caused by a perturbation. This study applied DN analysis, community identification and Reactome pathway analysis of the DN communities, and identified communities with highly statistically significant enrichment. We analyzed the DNs of two gene expression datasets where a perturbation was applied: (i) Saliva dataset (PPSV23 vaccination as perturbation; 24 time points), (ii) Primary B-Cells dataset (*ex-vivo* Rituximab drug treatment as perturbation; 6 time points). In summary, our results from the saliva DN revealed pathway activation in immunological and inflammatory responses. In the B cell DN, significant pathways were activated in the regulation of transcription, immune cell survival, activation and differentiation, and inflammatory response. By using the DN method on two separate data sets and comparing our results to known mechanisms of action and target pathways, we can assess the approach’s strengths and limitations. These are discussed further below.

### 3.1. PPSV23 Pathway Activations Following Perturbation in Saliva

*Streptococcus Pneumoniae*’s virulence and associated host immunity have been extensively studied [42]. The PPSV23 is an inactivated vaccine that uses purified capsular polysaccharides, and is typically administered to older adults (65+) and susceptible younger individuals [43, 44, 45, 46]. In our assessment we focused on the vaccine’s potential pathways of action. Our initial saliva investigation in PPSV23 established that an immune response to the vaccination can be detected utilizing non-invasive saliva monitoring at the molecular level [15]. Since aggregate saliva was sampled, we expected that multiple the multiple immune cells contained therein are involved in the observed patterns and associated immune responses. Based on our previous findings and general vaccine responses, we anticipated the activation of pathways involved with antigen presentation and processing, regulation of IgM and B/T cells, Lymphoid cells, MHC molecules, and phagocytosis. We also expected the activation of pathways of general immune response to stimuli or inflammation. To evaluate whether the DN method was as effective as previous studies, we focused on identifying the specific pathways involved.

In our results, a number of expected pathways emerged. These included pathways associated with antigen presentation and processing, Class I MHC mediated antigen processing and presentation, and ER-phagocytosis, and pathways governing the immunoregulation of interactions between Lymphoid and non-Lymphoid cells [38]. Further results indicative of the participation of immune cells, included the CLEC inflammasome pathway in C4. This pathway is associated with enabling host immune system to mount a fungal/bacterial defense using T-Helper 17 cells (TH17) [47, 48]. Interferon signaling, cytokine signaling, immune/adaptive immune and interleukin stimulation and signaling are all part of a generalized immune response [49]. We found these more general pathways in the pathway enrichment analysis of C0, C1, C2, C9, and C10. Interferon signaling is crucial in antiviral defense, cell regulation and growth, and immune response modulation [50]. Our Reactome pathway analysis results are consistent with the results of our saliva multi-omics study [15], which observed that vaccination activates various immune response and regulation pathways, which are also identified in our present results, including ER-Phagosome pathway, Interferon alpha/beta and gamma signaling, cytokine signaling, and MHC antigen presentation.

### 3.2. Rituximab Pathway Activations Following Perturbation in Primary B Cells

Regarding our primary B cell results, previous work [51] has established both the biological pathways and the mechanisms of action associated with Rituximab. These previous studies have demonstrated Rituximab’s ability to cause antibody-dependent and complement-dependent cellular cytotoxicity, growth inhibition and apoptosis, and regulation of the cell cycle. We also expected to observe Rituximab regulations of the B cell receptor (BCR) based on prior research. Particularly significant among our findings was the enrichment of the nuclear factor *κ*B (NF-*κ*B) pathways. According to Jazirehi et al. (2005) [23] and Bonavida (2005, 2007) [52, 53], treating NHL B cell lines with Rituximab inhibits NF-*κ*B’s signaling pathways by up-regulating RKIP and Raf-1 kinase inhibitors. RKIP has been found to antagonize signal transduction pathways that mediate the NF-*κ*B activation [54].

Following NF-*κ*B’s down-regulation due to RKIP’s up-regulation, the Bclxl expression is also down-regulated. As a result, tumor cells become more chemosensitive. Rituximab also decreased the activity of NF-*κ*B-inducing kinase, IkB kinase, and IkB-a phosphorylation. Finally, the introduction of Rituximab also decreased the activity of the IKK kinase and NF-*κ*B binding to DNA from 3 to 6 hours after treatment [23].

Among the more general enriched pathways observed are signaling pathways that play a role in the molecular mechanisms of chemosensitization, which are also impacted by Rituximab. In line with those effects, we anticipate impacts in the MAPK signaling pathway, the interleukin cytokine regulatory loop, and the Bcl-2 expression. Concerning the expression of genes involved in the healing process, research has uncovered Rituximab’s role in affecting pathways associated with immunoglobulin production, chemotaxis, immune response, cell development, and wound healing. Rituximab can also increase existing drug-induced apoptosis [51].

In our community of C4, for example, our Reactome analysis found 5 NF-*κ*B related pathways with FDR < 0.05. Of these 5 pathways, one is shown in Table 1; the remaining are displayed in the comprehensive table in the “Results/Bcell_results/reactome_analysis” folder on Zenodo. Alongside these NF-*κ*B pathways in C4 is the BCR pathway. Our results indicate that the C4 community response is highly relevant because of the activation of both NF-*κ*B and BCR pathways.

Our C2 community appears to be involved with the metabolism of proteins and cellular responses to external stimuli. Rituximab targets the CD20 B cell transmembrane protein that is involved in B-cell development, activation and proliferation [51]. The C2 community captures cell development pathways which were included in our expectations of more generalized responses.

We also observed relevant responses in other communities. For example, the C8 community showed activity in the RAF/MAP kinase cascade pathway. In a similar fashion, C10 demonstrated CD22 mediated BCR regulation, classical antibody-mediated complement activation, FCGR activation, antigen activation of the BCR, and initial complement triggering, etc. The pathways that emerged in our results are thus consistent and highly overlap with established pathways from previous studies. This suggests the effectiveness of our DN method.

### 3.3. Perturbation Induces Temporal Responses

Communities aid in defining the genes’ collective behavior, and observing the collective behavior of communities in the entire network can clarify relative trends between these collective behaviors. The generated heatmaps for each community depicted gene regulation for individual time points, and also displayed trends over time within the identified communities. The trends we observed in our saliva data were consistent with a time-dependent regulation. The results suggest a sequence of communities activations (up- and down-regulation) at individual timepoints, indicative of sequential immune system responses due to the PPSV23 vaccination. In the primary B cell data were less clear, as fewer time points were monitored, and also the network was more densely connected. The B-cell heatmaps still indicate overall trends associated with Rituximab activation (both up- and down-regulation) within the first 7 hrs of the treatment. Our future work will focus on the possibility of establishing a causal chained signaling response, and associated pathways across these communities.

### 3.4. Conclusion

Our analysis confirmed the applicability of a DN approach in evaluating time course RNA-seq data. Specifically, the DN method showed results in the saliva experiment data consistent with our previous work on profiling PPSV23 vaccination responses [15]. For the primary B cell responses to Rituximab, the DN has found the same signaling pathway as numerous prior experimental results, thus helping with our validation from a computational perspective. The DN approach complements prior studies by offering a systems-level network perspective of aggregate temporal changes due to drug activation. In future work we plan to address the identification of sequential activation of network communities, as well determining directionality/causality in such activations.

## 4. Limitations of the Study

Though our analysis identified multiple pathways relevant to Rituximab activation in the primary B cell data, heatmaps trends were not as distinct as those obtained from the saliva experiment, with weaker and less structured signals from the B cells. One major factor that may be contributing to these somewhat diffuse responses could be the nature of the *ex-vivo* experiment from which the data were obtained. Isolation in an *ex vivo* environment curtails interactions between Rituximab and aspects of the immune system that are difficult to measure efficiently using existing methods. This is in contrast to *in vivo* settings in which B cells have the ability to interact with those immune system factors. A key difference between the B cell data and the saliva data is that the latter were obtained under *in vivo* conditions, and thus reflect biological reality. In general, *ex vivo* experimental data are less accurate in summarizing the effects than those of *in vivo* experiments.

In addition, while the saliva experiment covered 24 time points over as many hours, the B cell experiment covered 6 time points over 15 hours. The sampling for the B cell dataset was both less frequent compared to the saliva dataset and unevenly spaced, thereby not accounting for the longer intervals during which no data were recorded.

Regarding the results from the saliva experiment, the data describes a bulk behavior from a tissue containing a mixture of different cell types instead of a single cell type. In principle, single cell RNA-seq data may provide better representation of dynamics and pathways involved in the response compared to a bulk RNA-seq dataset, and elucidate the temporal behavior of different individual cells involved (though currently such studies would also be limited to a pseudotime approach as each cell is only sampled once).

Finally, our DN method did not use the time information embedded in our time series dataset. Correlation based approached do not respect time ordering. Other methods like the Bayesian method and the causality inference method and may be helpful in determining the directionality of the edges in the gene network. As discussed above, we anticipate future work utilizing such methods may enable us to provide deeper information on the causal rewiring of the gene signaling network.

## Supporting information

Key Resources Table

## 5. Author Contributions

Conceptualization, S.X, C.P. and G.I.M; Methodology, S.X, C.P. and G.I.M.; Software, S.X. and G.I.M.; Investigation, S.X., L.R.K.R., M.Z., J.H, C.P. and G.I.M.; Visualization, S.X.; Resources, S.X., L.R.K.R., C.P. and G.I.M.; Writing – Original Draft, S.X., C.P. and G.I.M.; Writing – Review & Editing, All authors; Funding Acquisition, C.P. and G.I.M.; Supervision, C.P. and G.I.M.

## 6. Acknowledgements

This work was supported by the Translational Research Institute for Space Health through NASA Cooperative Agreement NNX16AO69A (project T0412). C.P also acknowledges support from NIH/NIGMS (R01GM122085).

## 10. STAR Methods

### RESOURCE AVAILABILITY

#### Lead contact

- Further information and requests for resources should be directed to and will be fulfilled by the lead contact, Shuyue Xue

#### Materials availability

- This study did not generate new unique reagents.

#### Data and code availability

- This paper analyzes existing, publicly available data. These accession numbers for the datasets are listed in the key resources table.
- Mapped RNA-seq data have been deposited at Zenodo and are publicly available as of the date of publication. DOIs are listed in the key resources table.
- All original code has been deposited on Zenodo as of the date of publication. DOI is listed in the key resources table.
- All results files have been deposited on Zenodo as of the date of publication together with the code files. These files are referred to as Online Data Files (ODFs) in the manuscript. DOI is listed in the key resources table.
- Any additional information required to reanalyze the data reported in this paper is available from the lead contact upon request.

### METHOD DETAILS

#### Data acquisition

Data for this investigation were obtained from Gene Expression Omnibus (GEO) for two time series studies using RNA-seq experiments, on Saliva (accession GSE108664) and Rituximab (GSE100441). Both sets of data are further described below. The raw RNA-seq data were mapped using Kallisto [58], with bootstrap sample parameter, `-b`, was set to 100. GENCODE[36] v28 transcripts and genome built GRCh38.p12 were used for annotation. We used Sleuth [59](with DESeq[60] adjustment of Transcripts per Million) to compile results across timepoints.

The saliva dataset was obtained from our previous study of immune responses to the PPSV23 vaccine (GSE108664) [15]. In this study, hourly saliva samples were collected from a healthy individual over two 24 hour periods and profiled with RNA-seq every hour. The first 24 hour period provides a baseline RNA expression dataset, which we call *untreated* data. In the second 24 hour period, the same individual was monitored after receiving the PPSV23 vaccine. Saliva samples were again collected hourly over 24 hours and profiled by RNA-seq. This second step yielded the RNA expression dataset after the PPSV23 vaccination. We call these data the *treated* dataset. Both treated and untreated datasets have 24 time points of 84,647 possible expression signals using GENCODE annotation [36].

The perturbation in the primary B cell experiment was Rituximab, a monoclonal antibody drug used in the treatment of different types of lymphomas and leukemias. The experimental study (data from GSE100441) began by culturing in parallel primary B cells with and without Rituximab. During the 15 hours of Rituximab treatment, the treated and untreated primary B cells were both sampled at the same 6 time points simultaneously and profiled by RNA-seq. The untreated group provided a baseline, which we call untreated data, whereas the treated experiment produced the treated dataset. Since this study included a replicated experiment, each of the first and second duplicates was processed to generate a separate network.

#### Data Preprocessing

For quality control, we pre-processed the experimental data and filtered sparse gene signals right after importing the published data files. We coded all the data analysis in Python in this study. Using Python’s pandas package [61, 62], we checked for missing values for each gene’s expression, removing duplicate records and eliminating genes with constant values across all the 24 time points for the saliva dataset (6 time points for the B cell datasets).

We replaced missing signals with zero and also set values less than 1 to 1. Genes with zero variance in their time series were excluded in our analysis. Moreover, we considered a gene signal as sparse and removed it if its time series had missing values for more than 1/8 of the time points. The same quality control procedure was used for both the saliva and primary B cell datasets.

#### Gene Selection

After quality control, we further processed the data to pre-screen and identify a pool of candidate genes that showed a significant response to the perturbations (vaccine in the saliva and Rituximab in the B cell). We selected genes that are significantly expressed in both untreated and treated cases. For each of these genes, we calculated the the time averaged relative difference between treated and untreated normalized intensities, Δ_*T U*_ :

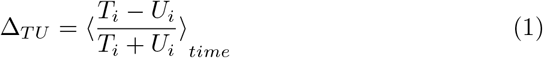

This calculation yielded a Δ_*T U*_ distribution curve, from which we computed the lower and upper quartiles. Genes were assumed to either be down-regulated or upregulated if their Δ_*T U*_ s were within the bottom 25% or top 25% of the Δ_*T U*_ distribution respectively. The Python Pandas package was used for all the above computations [61, 62].

#### Correlation Networks Construction

After gene selection, we calculated their pairwise Pearson correlation coefficients and built the co-expression networks. Genes were represented as nodes and were joined by edges if there was a non-zero correlation between them. We used the correlation coefficient as a weight for each edge. In the layout representation of the networks, the node-node distance reflects their correlation coefficients. Two genes are nearby if they have a high positive correlation. They are far apart when they have a low positive correlation or remote if negatively correlated. We used Python’s open source Networkx package [63] for network visualization and calculation of the network metrics.

To keep only the most significant signals, we kept edges only in the 99.5% quantile of the correlation distribution. This process removed many links on the network, detaching some nodes from others. We excluded those isolated nodes from the network. For the saliva data, we built one treated and one untreated network. Since we have data from two repeated experiments for B-cells, we built two networks for the Rituximab treatment and two networks for the untreated control. Then, we took the intersections between the two networks corresponding to the repeats to obtain a single Rituximab-treated network and one single control network.

#### Differential Networks Construction

We defined the DN as the control network subtracted from the treated networks both for the saliva and B cell case. The subtraction removed edges existing in both treated and untreated networks and preserved those only present in the treated network. This operation generates some isolated nodes, that were discarded.

We analyzed the DN’s structure using modularity [25, 26], as complex biological networks usually display a high degree of modularity [29]. Specifically, we employed the Louvain community detection method [34], one of the best-known algorithms for its efficiency, to partition the entire DN into smaller clusters, also known as communities. The Louvain method has been integrated into a published Python package [64], and we used this existing implementation. This graph clustering algorithm is not deterministic, and can therefore result in different partitions for the same graph.

#### Enrichment Analysis and Heatmaps

We conducted Reactome Enrichment Analysis [35] on each community to identify over-represented biological pathways within each community. Reactome Enrichment Analysis was performed using the Python package PyIOmica [37]. As the majority of communities with low number of genes yield no enrichment, we focused on results for communities with 8 or more genes. Furthermore, we plotted the heatmaps for individual communities to visualize gene expression levels as time series, exploring the communities’ collective behavior as a function of time. On the heatmaps, we plotted differences in gene expression relative to time 0 (first time point). We computed these values by subtracting each gene’s individual time points from time 0. Each time series in our heatmaps was normalized using the Euclidean norm. For dendrogram clustering of rows (genes), we applied the complete-linkage method (Farthest Point Algorithm) [55, 56].

#### Results formatting and visualization

We stored the DN nodes and edges, communities, and pathway enrichment analyses into spreadsheets that are provided in the ODFs both for the saliva and B cell data. Using Mathematica [65], we visualized the saliva and B cell DNs with their major connected components and communities.

### QUANTIFICATION AND STATISTICAL ANALYSIS

The data were quantified as discussed in the Method Details section above. Statistical considerations included: (i) selection of a correlation cutoff in network edge construction, based on a 0.995 quantile of the correlation distribution. (ii) An FDR < 0.05 cutoff was used to assess statistical significance of Reactome pathway enrichment analysis, generate through the use of the Reactome API in PyIOmica[35, 37].

