## Supplementary material for "Differential Network Analysis of Longitudinal Gene Expression in Response to Perturbations": Key Resources Table

| **REAGENT or RESOURCE** | **SOURCE** | **IDENTIFIER** |
| --- | --- | --- |
| **Deposited Data** | | |
| Saliva mRNA-sequencing | Gene Expression Omnibus (GEO) | GSE108664 |
| Rituximab Treatment in Primary B Cells mRNA-sequencing | Gene Expression Omnibus (GEO) | GSE100441 |
| Online Data Files (ODFs; Code & Results from this paper) | This paper | doi: 10.5281/zenodo.5519804 |
| **Software and Algorithms** | | |
| Louvain community detection algorithm | (Blondel et al., 2008) doi:10.1088/1742-5468/2008/10/p10008 | doi: 10.1088/1742-5468/2008/10/p10008 |
| Scikit-network: Graph Analysis package in Python 0.24.0 | https://jmlr.org/papers/v21/20-412.html | https://scikit-network.readthedocs.io/en/latest/ |
| PyIOmica 1.3.1 | (Domanksy et al., 2020), doi:[10.1093/bioinformatics/btz896](https://doi.org/10.1093/bioinformatics/btz896) | https://pyiomica.readthedocs.io/en/latest/ |
| Mathematica 12.0.0 | Wolfram Research | https://www.wolfram.com/mathematica |
| Pandas package in Python 1.1.3 | (The Pandas Development Team, 2020) doi:10.5281/zenodo.3509134 | https://pandas.pydata.org/ |
| Networkx package in Python 2.5 | (Hagberge et al., 2008) | https://networkx.org/ |
